# Clinical Impact and Cost-Effectiveness of a 176-Condition Expanded Carrier Screen

**DOI:** 10.1101/372334

**Authors:** Kyle A. Beauchamp, Katherine A. Johansen Taber, Dale Muzzey

## Abstract

**Purpose:** Carrier screening identifies couples at high risk for conceiving offspring affected with serious heritable conditions. Minimal screening guidelines mandate testing for cystic fibrosis and spinal muscular atrophy, but expanded carrier screening (ECS) assesses reproductive risk for hundreds of conditions simultaneously. Although medical societies consider ECS an acceptable practice, the health economics of ECS remain incompletely characterized.

**Methods:** The clinical impact and cost-effectiveness of a 176-condition ECS panel were investigated using a decision-tree model comparing minimal screening and ECS in a preconception setting. Carrier rates from >50,000 patients informed disease-incidence estimates, while cost and life-years-lost data were aggregated from the literature and a cost-of-care database. Model robustness was evaluated using one-way and probabilistic sensitivity analyses.

**Results:** For every 100,000 pregnancies, 300 are predicted to be affected by ECS-panel conditions, which, on average, individually incur $1,300,000 in lifetime costs and increase mortality by 26 undiscounted life-years on average. Relative to minimal screening, ECS reduces the affected-birth rate and is cost-effective (i.e., <$50,000 incremental cost per life-year), findings robust to reasonable model-parameter perturbation.

**Conclusion:** ECS is predicted to reduce the population burden of Mendelian disease in a cost-effective manner compared to many other common medical interventions.

## Introduction

Collectively, Mendelian diseases account for approximately 20% of infant mortality and 18% of infant hospitalizations^1^. To address this significant health concern, the American College of Obstetricians and Gynecologists (ACOG)^2,3^, the American College of Medical Genetics and Genomics (ACMG)^4^, and other medical societies have recommended carrier screening for select prevalent conditions. ACOG recommends cystic fibrosis (CF) and spinal muscular atrophy (SMA) screening for all women considering pregnancy (or already pregnant)^3^, as well as additional screening based on family history and ethnicity.

Expanded carrier screening (ECS), often performed using next generation sequencing (NGS), screens tens to hundreds of conditions^5,6^. ECS disease panels can be designed using systematic principles (e.g., severity)^7,8^, ECS testing can be performed with high sensitivity and specificity^9^, and screening results lead to measurable changes in reproductive decision-making^10–12^. Large-scale retrospective analyses have shown that traditional carrier screening may fail to detect many pregnancies affected with Mendelian disease, often disproportionately impacting select ethnic groups^13^.

Though examined for individual conditions^14–16^, cost-effectiveness of ECS is not fully characterized. One study found NGS-based ECS with a 14-condition panel to be cost-effective^5^. Others, though not directly addressing ECS, examined the cost-effectiveness of treatments: due to high prices, recently approved drugs treating ECS conditions (e.g., CF^17^ and SMA^15^) may not be cost-effective.

Two recent actions from medical societies could impact ECS utilization. ACOG has recognized ECS as an acceptable screening strategy for reproductive risk management^2^, and the American Medical Association CPT Editorial Panel has approved a CPT code (81X43, effective Jan. 2019) for panethnic sequencing-based ECS panels with ≥15 genes^18^. With these actions potentially leading to near-term increased use of ECS, now is an opportune time to evaluate the impact of ECS on the health system.

Cost-effectiveness analysis^5,19–21^ evaluates the cost and clinical benefits of medical interventions. It condenses each comparison among interventions into a single number (incremental cost-effectiveness ratio; ICER) that summarizes the cost per unit of clinical outcome (e.g., dollars per life-year). Here we model the clinical impact and cost-effectiveness of ECS, building upon previous work (e.g., ref.^5^) in several ways. First, we consider a large panel with 176 conditions. Second, we aggregate disease cost and life-years-lost data from both literature and cost-of-care databases. Third, we model disease incidence using a cohort of more than 50,000 patients. Finally, we use a decision model to compare the cost-effectiveness of ECS relative to a minimal screening protocol or no screening.

## Materials and Methods

### Decision Tree

Fig. 1 summarizes the carrier-screening workflows compared in this work. The model was chosen as the most parsimonious protocol that captures the key aspects of carrier screening. First, couples may be at risk or not at risk for a panel condition. Second, we assume that the detection rate for each condition is 100% (see Discussion), so that couples deemed “not at risk” have zero chance of having a child affected with a screened condition (see Discussion). Third, at-risk couples (ARCs; defined below) have a fixed probability of choosing a reproductive intervention that is assumed to avert any possibility of a child affected by a screened condition; this fixed probability (76%) is estimated from survey-based clinical utility studies of ARCs^10,11^. Here we focus on preconception screening (see Discussion).

**Figure 1.**
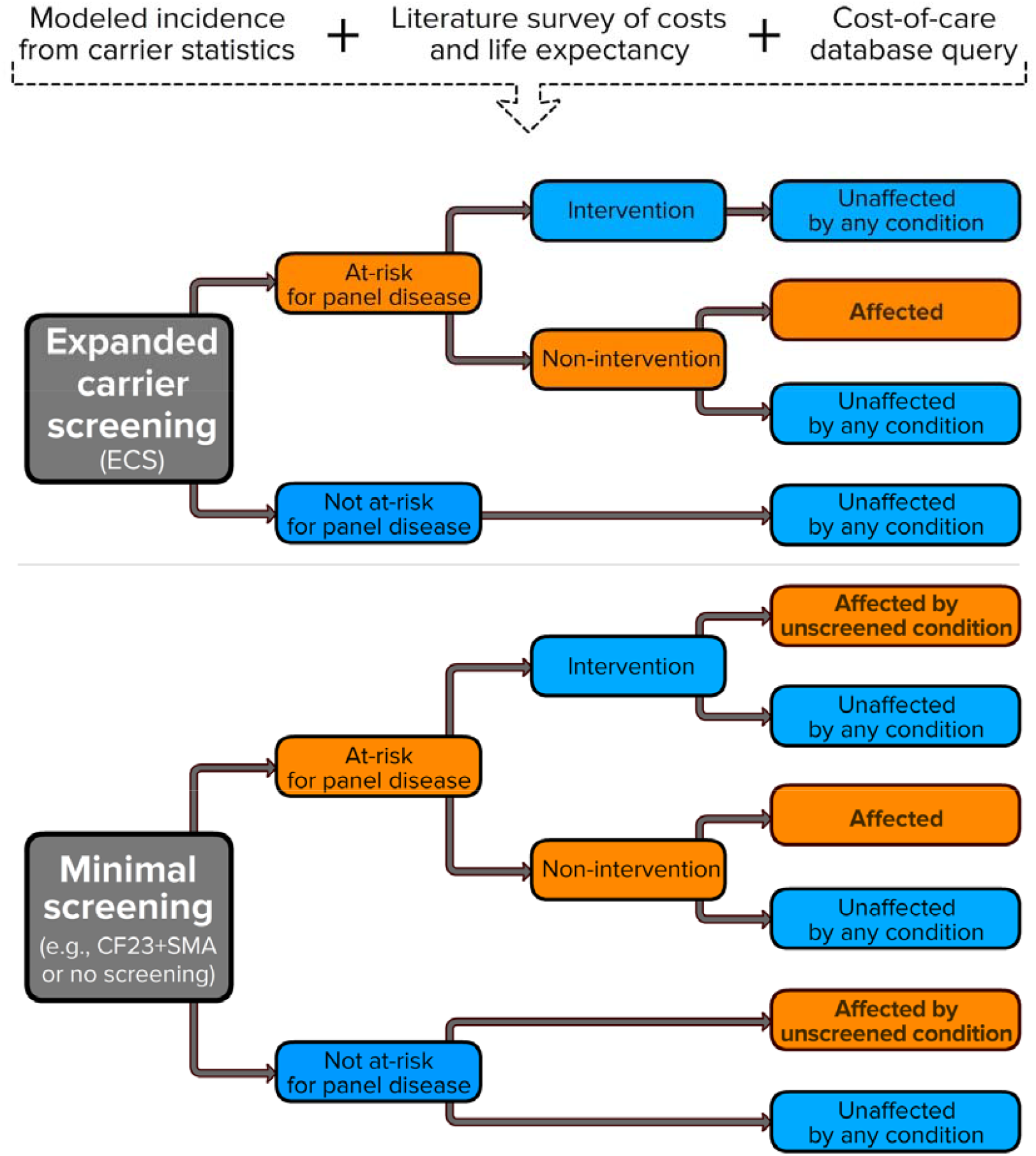
Decision-tree model. Top): analyses in this work combined three data sources to construct a decision-tree model. Bottom): the decision-tree model considered in this work—one arm for expanded carrier screening (ECS) and another for minimal screening (CF23+SMA screening or no screening)—depicts the risk-status, intervention choice (where applicable), and pregnancy outcome from left to right. Risk-mitigating and unaffected nodes are blue, whereas risk-increasing, risk-maintaining, and affected nodes are orange. The decision tree, like the rest of the modeling described herein, assumes that ECS tests for the totality of diseases; therefore, “Unaffected by any condition” means that the pregnancy is not affected for a condition screened on the ECS panel, and “Affected by unscreened condition” means a pregnancy is affected by a condition on the ECS panel beyond CF23+SMA. In the analyses herein, six variants of these subtrees are considered.

### Institutional Review Board Approval

This study was reviewed and designated as exempt by Western Institutional Review Board.

### Disease List

This study examines diseases on the 176-condition Foresight ™ Universal Panel^9^ (see Supplementary Table S1), currently used for ECS at Counsyl.

### Data Sources: Overview

Our analyses utilize per-disease estimates of incidences, life-years lost, and disease-treatment costs, which we collected from three sources (Fig. 1; top). First, aggregated carrier statistics were used to model fetal-disease incidences (see “Incidence Modeling”). Second, a literature survey of disease-cost and life-expectancy data was performed (see “Literature Survey”). Third, we acquired data on aggregated health outcomes and expenses for a longitudinal cohort of millions of patients (Truven Health Analytics; see “Cost-of-care Database Survey” and Supplementary Table S4). Supplementary Tables S2 and S3 contain the value and provenance of quantities used in this work.

Disease cost estimates were converted to 2018 dollars assuming a medical-care inflation rate of 3.6%^22^ per year. When from a literature source, life-years and lifetime-cost data were assumed to be present-value discounted appropriately. When such parameters were modeled herein (e.g., from lifespan data; see Supplementary Methods), life-years lost and lifetime costs were present-value adjusted assuming a discount rate of 3%^23^.

### Incidence Modeling

Decision-tree models were based on US disease-incidence estimates. Because affected persons for each condition are rare, we modeled disease incidence using the inheritance patterns (autosomal recessive or X-linked) and carrier frequencies estimated from 53,163 patients screened by Counsyl using the 176-condition Universal panel. This “modeled fetal disease risk” (MFDR) approach, described previously^7,13^, predicted the frequency of affected conceptuses. To reduce bias, the cohort excluded patients with fertility issues or family history of disease.

To estimate the rate of at-risk couples (ARCs) in which both partners are carriers for an autosomal-recessive condition or the mother is a carrier for an X-linked condition, we note that for both autosomal recessive and X-linked conditions, the ratio of ARC rate to MFDR is 4, so we define the ARC rate as 4 * MFDR. For diseases with complex inheritance (e.g., fragile X syndrome), this approximation simplifies the decision analysis. Panel-wide MFDR and ARCs were treated as additive across the panel’s constituent diseases, consistent with previously described methods^7^. Calculations assumed intra-ethnicity coupling and were reweighted to match the ethnic makeup of the general US population^24^, as described previously (Supp. Table S1 in ref. [^7^]).

### Literature Survey

We performed a literature survey to aggregate the following quantities for each condition: lifetime disease cost, years 0-3 (post-birth) disease cost, years 0-1 (post-birth) disease cost, and life-years lost. Because many panel conditions have been studied most carefully by nonprofit and governmental agencies, we included scientific publications, government resources (e.g., from the National Institutes of Health), reports from disease support groups, and clinical resources for patients and genetic counselors as acceptable literature. For cost estimates of several orphan drugs (e.g., nusinersen / Spinraza™) that recently obtained FDA approval, we used FDA indication documents, reports from published news sources (e.g., Forbes), and reports from trade organizations (e.g., America’s Health Insurance Plans; AHIP). For 51 conditions, we also used lifetime-cost estimates from our previous analysis of treatment costs^25^.

To estimate the cost of follow-up reproductive care for the ARCs who intervene to avoid an affected birth, we integrated the probabilities and costs of pre-implantation genetic diagnosis (PGD), in vitro fertilization (IVF), prenatal diagnosis (PD), and termination from Table 1 in ref. [^5^]; each reproductive intervention was estimated to cost $14,312.04, primarily driven by the cost of IVF.

**Table 1.** Disease frequency and cost analysis. U.S.-weighted frequency, cost per affected birth, and cost per unscreened birth weighted by frequency are tabulated for the ten most common diseases. Cost figures are shown for the lifetime, 0-1yr and 0-3yr time periods, where year 0 is birth.

| Disease | Frequency<br>(1 in X) | Lifetime<br>Cost | Years 0-3<br>Cost | Years 0-1<br>Cost | Weighted<br>Lifetime Cost | Weighted<br>Years 0-3 Cost | Weighted<br>Years 0-1 Cost |
| --- | --- | --- | --- | --- | --- | --- | --- |
| Hb beta chain-related<br>hemoglobinopathy | 2,174 | \$632,613 | \$44,147 | \$16,663 | \$291 | \$20 | \$8 |
| cystic fibrosis | 2,511 | \$3,083,907 | \$95,910 | \$20,957 | \$1,228 | \$38 | \$8 |
| fragile X syndrome | 2,939 | \$974,614 | \$24,109 | \$4,259 | \$332 | \$8 | \$1 |
| dystrophinopathy<br>(including<br>Duchenne/Becker<br>muscular dystrophy) | 3,571 | \$1,507,227 | \$54,670 | \$7,135 | \$422 | \$15 | \$2 |
| GJB2-related DFNB1<br>nonsyndromic hearing<br>loss and deafness | 6,191 | \$12,238 | \$8,556 | \$1,529 | \$2 | \$1 | \$0 |
| phenylalanine<br>hydroxylase<br>deficiency | 8,683 | \$187,286 | \$294,830 | \$101,285 | \$22 | \$34 | \$12 |
| congenital adrenal<br>hyperplasia | 10,230 | \$170,821 | \$71,873 | \$30,314 | \$17 | \$7 | \$3 |
| spinal muscular<br>atrophy | 10,879 | \$2,546,777 | \$1,466,588 | \$750,000 | \$234 | \$135 | \$69 |
| Smith-Lemli-Opitz<br>syndrome | 12,243 | \$92,940 | \$209,429 | \$131,259 | \$8 | \$17 | \$11 |
| Fabry disease | 12,861 | \$5,115,951 | \$7,776 | \$0 | \$398 | \$1 | \$0 |

### Cost-of-care Database Survey

Short-term disease cost data were queried from a commercial database (Marketscan; Truven Health Analytics), commonly used in health-economics studies (see, e.g.,^26–28^). Twenty-five conditions were selected based on having high incidence and specific International Classification of Diseases (ICD) coding. For each disease, ICD9 or ICD10 codes were chosen based on which code was sufficiently specific and which database (ICD9 or ICD10) had a larger cohort. Supplementary Table S4 lists all coding choices.

For each disease, a number of queries were performed (e.g., costs for commercial payers and Medicaid separately, costs at different ages, number of patients with a diagnosis, etc.); the complete output of the Truven Health Analytics queries is given in Supplementary Table S4. For the modeling herein, only the first-year-cost and three-year-cost numbers were used (cells C37 and C39 on each conditions’ tab in Table S4). To estimate the disease costs during the first N years of life (where N is either 1 or 3), we did the following. First, patients were included based on age and continuous enrollment during the appropriate time window. Second, patients were labeled as being diagnosed or not diagnosed based on whether they received an appropriate diagnosis code at some point in the portion of their life chronicled in the Truven Health Analytics database. Finally, for both commercial payers (“com”) and Medicaid (“med”) separately, the annual average costs of the diagnosis cohort (“μ_dx”) and an appropriately matched (based on age, eligibility, time window, payer) non-diagnosis cohort (“μ_nodx”) were calculated. The single and final disease cost number reported for each disease enforced nonnegativity (disease_cost > 0) and, where possible, prioritized commercial payers (versus Medicaid), as indicated below:

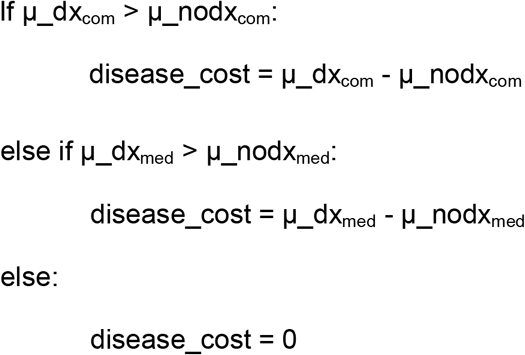

### Missing-value Estimation

Because certain values were missing due to incomplete data, we estimated the remaining values. For lifetime disease costs, we used the following rules: If available, use a literature estimate of the lifetime cost; otherwise, discount and accumulate the annual cost for each year in the life expectancy; if the annual cost or life expectancy is unknown, use the cost of the first three years of life; if the cost of the first three years of life is unknown, discount and accumulate three years of annual costs. For values unaddressed by these rules, we assigned the value to be the median of other available corresponding values. For example, the unavailable first-year cost for glycogen storage disease type Ia is estimated using the median first-year cost among all diseases for which values are available. Our robustness analysis (see “Sensitivity analysis”) measured the impact on model predictions of variations in these median missing-value estimates. Finally, in cases with multiple data points for a single cost parameter, we used the largest cost (due to medical inflation and new treatments, a recent estimate was generally selected).

### Screening modes

Our analyses examined screening modes selected to represent the different theoretical and practical ways in which carrier screening could be adopted and evaluated:

- **“No Screening”**. Couples are not screened for any panel condition.
- **“Population Impact: CF23+SMA”**. Couples are screened for the 23 ACMG-recommended CF variants^29^ and SMA. 100% of ARCs intervene to avoid an affected birth. Couples may still be at risk of conditions beyond CF23 and SMA.
- **“Population Impact: ECS”**. Couples are screened for 176 conditions. All ARCs intervene to avoid an affected birth.
- **“76% Intervention CF23+SMA”**. Couples undergo CF23+SMA screening, but 76% of ARCs intervene to avoid affected births, consistent with findings in (^10^).
- **“76% Intervention ECS”**. Couples undergo ECS in which 76% of ARCs intervene to avoid affected births.

The two “Population Impact” models were included to isolate the population impact of panel diseases (ECS or CF23+SMA) irrespective of the intervention behavior of ARCs; these results are equivalent to the behavior of a hypothetical cohort in which 100% of ARCs intervene to avoid an affected birth. Overall, we considered three primary screening modes (No screening, CF23+SMA screening, and ECS) but examined the respective population impacts of the underlying diseases and the screening interventions that ARCs pursue.

### Sensitivity Analysis

We assessed model sensitivity in two ways^19,20^: (1) one-way sensitivity analysis, in which all but one variables were held at their base values and the remaining value was set to levels that capture realistic bounds on the variable,and (2) probabilistic sensitivity analysis, whereby all variables were drawn from realistic prior distributions and outcome metrics are assessed across the ensemble of sampled parameter vectors. Model parameters, base values, ranges, and prior distributions are summarized in Supplementary Table S2, and described in the Supplementary Methods.

### Software

Analyses were performed using python (software versions in Supplementary Table S6).

## Results

### Disease Incidence and Cost Estimates

For each disease on a 176-condition ECS, we estimated incidence and treatment costs based on screening results from 53,163 patients, a detailed literature survey of ~80 published resources, and query results from a cost-of-care database (See Methods); results for the panel’s 10 most frequent conditions are in Table 1 (full results in Supplementary Tables S3 and S5). Disease-incidence rates span multiple orders of magnitude: common diseases like beta chain hemoglobinopathies, CF, Duchenne/Becker muscular dystrophy, and fragile X syndrome together affect more than 100 per 100,000 pregnancies, whereas the panel’s 150 least-common diseases affect approximately 40 per 100,000 pregnancies. Lifetime cost estimates are also highly variable, with several diseases exceeding $1,000,000 per affected individual. Early costs are skewed as well, due to some diseases having onset in infancy (e.g., SMA) and others in adolescence/adulthood (e.g., Fabry disease).

### Clinical and Economic Impacts of Screening

We built a decision-tree model (Figure 1; see Methods) to assess the population-level impact of ECS diseases in terms of both clinical outcomes (e.g., number of ARCs, affected births averted, and life-years lost) and economic factors (e.g., spending over first year, first three years, and lifetime) (Table 2).

**Table 2.** Model results per single-birth couple. Costs exclude ECS screening costs, which are considered separately in the ICER analysis (Fig. 2). All values are reported as per couple rates.

| Panel | At risk<br>Couples<br>Detected | Number of<br>reproductive<br>interventions | Affected<br>Births<br>Averted | Life-years<br>Gained | Life-years<br>Gained<br>(Undiscounted) | Lifetime<br>Cost<br>Averted | Three<br>Year<br>Cost<br>Averted | Year<br>One<br>Cost<br>Averted | Interventi<br>on Costs<br>Accrued |
| --- | --- | --- | --- | --- | --- | --- | --- | --- | --- |
| **Population**<br>**Impact: ECS** | 1.20E-02 | 1.20E-02 | 3.00E-03 | 1.74E-02 | 7.85E-02 | \$3,823.76 | \$466.69 | \$166.75 | \$171.99 |
| **Population**<br>**Impact:**<br>**CF23+SMA** | 1.49E-03 | 1.49E-03 | 3.72E-04 | 4.98E-03 | 1.89E-02 | \$1,097.90 | \$161.67 | \$74.81 | \$21.30 |
| **Population**<br>**Impact:**<br>**Difference** | 1.05E-02 | 1.05E-02 | 2.63E-03 | 1.24E-02 | 5.96E-02 | \$2,725.86 | \$305.02 | \$91.94 | \$150.69 |
| **76% Intervention**<br>**ECS** | 1.20E-02 | 9.16E-03 | 2.29E-03 | 1.33E-02 | 5.98E-02 | \$2,913.34 | \$355.57 | \$127.05 | \$131.04 |
| **76% Intervention**<br>**CF23+SMA** | 1.49E-03 | 1.13E-03 | 2.83E-04 | 3.80E-03 | 1.44E-02 | \$836.49 | \$123.18 | \$57.00 | \$16.23 |
| **76%**<br>**Intervention:**<br>**Difference** | 1.05E-02 | 8.02E-03 | 2.01E-03 | 9.46E-03 | 4.54E-02 | \$2,076.84 | \$232.40 | \$70.05 | \$114.81 |

In terms of clinical impact, our modeling predicted that 1.2% of couples will be at high risk of one of the 176 screened conditions, with 300 in 100,000 births affected (Table 2: “Population Impact:”). Due to the high mortality associated with ECS conditions, the per-birth rate of lost life-years was 0.017, or 0.078 without discounting (Table 2). In clinical practice where screening is performed, approximately 76% of patients would intervene^10,11^ to avoid an affected birth (via, e.g., in vitro fertilization with preimplantation genetic diagnosis, use of donor gamete, adoption, or decision to avoid pregnancy), meaning that ECS would avert 230 affected births per 100,000 pregnancies (Table 2: “76% Intervention ECS”). Finally, the model also estimated the incremental gain of ECS over minimal screening (“76% Intervention CF23+SMA” in Table 2), showing that ECS averts an additional 200 births per 100,000 pregnancies.

To explore the economic implications of ECS, we combined the estimates of incidence and disease cost to measure the incremental changes in medical spending associated with ECS panel conditions. Because different stakeholders (e.g., society, insurers) may prioritize different time horizons, we examined three periods: lifetime, years 0-3, and years 0-1, where year 0 is birth. This modeling focused entirely on disease-treatment costs and did not take into account the cost of screening or reproductive care (discussed later). Importantly, these cost figures represent mitigation of current financial outlays by insurers; they do not yet account for gain in health outcomes (e.g., increased life years) due to ECS, addressed below in the ICER analysis.

In the population-impact scenario, the avoided costs averaged over all births—the minority affected plus the majority unaffected—were approximately $3,800 in lifetime costs, $470 in years 0-3 costs, and $170 years 0-1 costs (Table 2). Among only affected births, the average lifetime cost per affected birth was $1,300,000. Considering a more realistic comparison wherein 76% of ARCs intervene to avoid an affected birth^10^ and CF23+SMA is used as a baseline, ECS still showed large incremental cost savings when averaged over all births: approximately $2,100 over a lifetime, $230 in years 0-3, and $70 in years 0-1.

Separately from the aggregation of future disease costs, we modeled the financial impact of reproductive care (PGD, IVF, PD, and termination) for those ARCs who intervene to avoid an affected pregnancy. These costs are aggregated in the column “Intervention Costs Accrued” in Table 2. Because ARCs are relatively rare (1.2% as modeled herein) and some choose not to pursue further intervention, the per-couple contribution of these costs is low ($170) when compared to the disease-treatment costs on the 3-year or lifetime horizons ($470 and $3800, respectively).

### Cost-effectiveness of ECS

We examined the cost-effectiveness of ECS (Fig. 2) by treating ECS price as a variable and plotting the incremental cost-effectiveness ratio (ICER) with or without modeled cost savings (for years 0-3) due to averted disease. Because insurer coverage for IVF expenses is highly variable, the modeling in Fig. 2 excludes such expenses (Fig. S1 shows the analysis including IVF expenses). For comparison, cost-effectiveness values obtained for hereditary cancer screening are also shown^20,31–33^.

**Figure 2.**
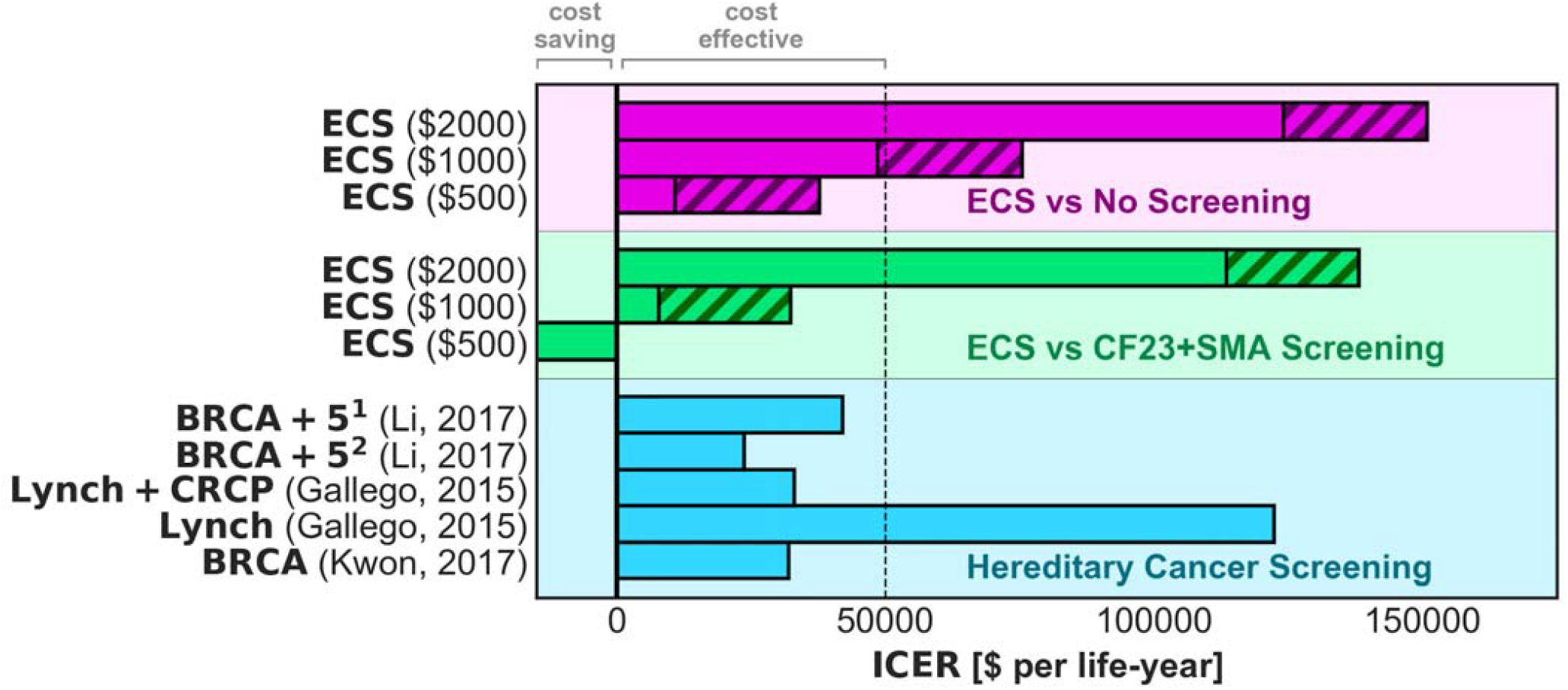
ECS cost-effectiveness. The incremental cost-effectiveness ratio (ICER) of “76% Intervention ECS” is compared to no screening (magenta) and CF23+SMA screening (green). Life-years gained are used as clinical outcomes of interest. ECS price is shown in parentheses. Solid bars indicate the ICER modeled with 3-year cost savings subtracted, while the level indicated by hatched bars indicates the ICER without such deduction (i.e., only accounting for screening cost). As a comparison, ICER values are shown for several inherited cancer interventions (blue). The common $50,000 per life-year cost-effectiveness threshold is shown as dashed lines; note, however, that thresholds as high as $100,000 have been proposed^38^. Superscripts (1) and (2) refer to multiple ICER estimates from the same study. In the green section, the third bar (“ECS ($500)”) is cost saving because after subtracting the price of CF23+SMA screening ($693.60), the cost per life-year is negative. The plot is truncated because negative ICER results are typically not interpreted quantitatively, as one alternative is superior to the other in terms of both cost and life-years saved. See Fig. S1 for a similar analysis that also includes IVF expenses.

Compared to no screening, “76% Intervention ECS” showed a cost-effectiveness near the common benchmark value of $50,000 per life-year^34^ (solid and hatched magenta bars in Figure 2). When cost-savings due to avoided disease are included in the analysis, the ICER of ECS improves substantially (solid magenta bars in Figure 2). For instance, if ECS were priced at $1,000 and hypothetically saves .01 life-years, the ICER is $100,000 per life-year; when taking into account savings of $400 in treatment costs, the ICER is ($1000 - $400) / .01 = $60,000 per life-year. Further, when costs are evaluated on a lifetime horizon (rather than 3-year horizon), ECS is cost-saving at prices up to $2,913 because the net averted costs are greater than the price.

Compared to CF23+SMA screening, “76% Intervention ECS” showed even greater cost-effectiveness (Fig. 2; lime). The price of CF23+SMA screening ($693.60) was estimated as the median price paid for CF23+SMA screening (CPT codes 81220, 81401) across US commercial laboratories, as reported under the Protecting Access to Medicare Act (PAMA)^35^. The improved cost-effectiveness of ECS in this comparison arose for two reasons. First, the price of CF23+SMA screening was large when compared to the ECS prices examined. Second, the majority of ECS clinical impact (affected births averted and life-years gained) stemmed from panel conditions beyond CF23 and SMA. Together then, the ICER of ECS relative to CF23+SMA was high because the improved clinical outcomes from ECS were achieved for a smaller marginal cost increase than the ICER calculations that used no screening as a baseline. Interestingly, at low price points (i.e., near CF23+SMA pricing of $693.60), ECS is highly cost-effective and incrementally cost-saving—that is, averted disease costs outweigh the incremental price difference between ECS and CF23+SMA screening.

### Sensitivity Analysis to Assess Robustness

To explore how susceptible the model’s conclusions were to changes in the underlying parameters, we assessed model robustness to both single-parameter variation (one-way sensitivity analysis) and multivariate-parameter uncertainty (probabilistic sensitivity analysis). In one-way sensitivity analysis (Fig. 3; bottom), we evaluated the impact of several model parameters: (1) the population ARC frequency, (2) the fraction of ARCs who alter their reproductive behavior, (3) the fraction of CF risk attributable to the 23 ACMG-recommend variants, and (4) the assumed lifetime cost value for conditions without a published value. As expected, the model was most sensitive to the population ARC frequency, which could vary greatly when applying ECS to high-risk (e.g., Ashkenazi Jewish), intermediate-risk (e.g., US population), and low-risk (e.g., East Asian) ethnicities. The model showed less pronounced sensitivity to the other model parameters.

**Figure 3.**
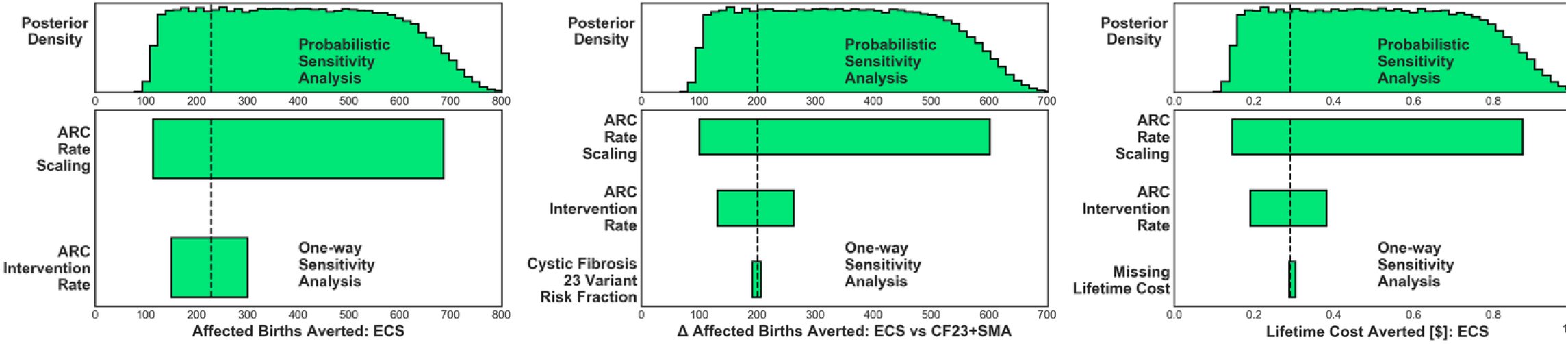
Sensitivity analysis. (bottom). One-way sensitivity analysis tornado plot shows sensitivity to model parameters for: Affected Births Averted by ECS, difference in Affected Births Averted between ECS and CF23+SMA screening, and lifetime costs averted by ECS. (top). Probabilistic sensitivity analysis shows the sensitivity of affected births averted due to multivariate parameter uncertainty. Results are per 100,000 couples.

The probabilistic sensitivity analysis largely reflected the range of values observed in the oneway sensitivity analysis when scaling the population ARC frequency (Fig. 3; top); this correspondence was unsurprising given the dominant contribution that this parameter showed in the one-way analyses. The probabilistic sensitivity analysis used the prior distributions enumerated in Table S2 and suggested that for every 100,000 births the median number of averted affected births was 398 (95% credible interval: 128-703).

## Discussion

Here, a decision tree model integrating incidence, clinical-impact, and cost data was used to reveal several new findings about ECS. First, the cost associated with screened diseases is large, even when averaged over a population of affected and unaffected births. Second, the clinical benefits of screening (affected births averted and life-years gained) are largely attributable to diseases beyond minimal guidelines-based screening (CF23+SMA). Third, the cost-effectiveness of ECS is favorable, in particular when averted disease costs are considered and when compared against CF23+SMA screening. We also observed that high-cost diseases tend to have high-priced, recently approved drugs (e.g., Spinraza) or long life expectancy with treatment (e.g., CF), suggesting that financial impact of Mendelian diseases may increase as new orphan drugs obtain approval.

Modeling carrier screening is complicated because of the choices involved in what to model, including whether screening occurs before or after conception, which interventions are chosen among different cohorts, and how well screening results correspond to affected births (e.g., due to screen sensitivity or disease-specific miscarriage rates). While more complex decision-tree models have attempted to capture several of these variations^5^, they required many uncertain parameters (e.g., rates of specific reproductive choices) and emphasized individual carrier status rather than the more clinically relevant (particularly for large panels) ARC status. In this work, we pursued the following approach: (1) model a simple ECS workflow that both captures the key clinical management steps of carrier screening and uses transparently ascertained or estimated parameter values, and (2) account for simplifying assumptions by assessing uncertainty in the model’s conclusions with a sensitivity analysis. Below we discuss some of the simplifying assumptions and their implications on our results.

While many couples undergo prenatal ECS, preconception screening was the focus of our model because it has well established utility and confers greater patient autonomy^2^. We expect similar results at either stage but with a reduced rate of reproductive interventions for prenatal screening^10^. Importantly, even if the intervention rate were to drop from its 76% preconception value to 50% in the prenatal period (as was observed in the post-conception cohort in^10^), our one-way sensitivity analysis demonstrated that ECS would still avert 150 births affected with severe disease and $2,000 in lifetime cost per birth. One policy implication of these results is that a payer obligated (by guidelines and medical policy) to provide CF23+SMA screening could cost-effectively increase member health by performing ECS, possibly with incremental cost savings over both short and long time horizons.

Our model generalizes the idea of reproductive intervention to avoid an affected birth—rather than modeling each intervention type individually—because patients make reproductive choices in accordance with their values. Thus, the distribution of particular reproductive interventions would depend on the cohort of interest and the testing timeline. These dependencies are mitigated by instead focusing on the clinical outcomes of interest (e.g., affected births averted), which may be more generalizable than the specific interventions chosen. Furthermore, from the economic perspective, expenditures associated with reproductive interventions such as IVF and PGD may accrue to different parties, depending on the couple’s insurance. In Table 2, we separately accounted for disease costs and reproductive-intervention costs, with the understanding that stakeholders assess these costs differently.

Regarding the correspondence between preconception carrier screening results and the risk of an affected birth, we assumed perfect sensitivity and specificity for screened diseases, and we did not take into account the impact of miscarriage. The assumption of perfect test accuracy—which we submit is justified given the >99% analytical sensitivity and specificity reported in the validation of the 176 condition ECS panel^9^—means that an affected birth can only result from either not screening for a particular disease or an ARC not pursuing intervention. Though miscarriage can be common for certain diseases (e.g., Smith-Lemli-Opitz syndrome^36^) and may impact cost estimates, we expect this impact to be minor because few diseases and variants yield higher miscarriage rates.

Although not the focus of this study, it is important to note that ECS provides benefits beyond the reduction of affected births and the increase in expected life-years. These include early education for conditions associated with intellectual disability (e.g., in fragile X syndrome^37^), early detection of impairment that can lead to intervention or treatment (e.g., cochlear implants to treat *GJS2*-related deafness), communication of risk to family, and faster diagnoses of rare disorders that could otherwise require a diagnostic odyssey. Importantly, many of these benefits accrue to couples who choose not to avoid an affected pregnancy. Though they are difficult to account for in a quantitative cost-effectiveness framework, these benefits are important to consider.

## Supplementary Information

Supplementary information is available online as a separate PDF.

